# Waterbird abundance and diversity along a stretch of Ganga River, India modified by construction of a barrage

**DOI:** 10.1101/2020.12.17.423209

**Authors:** Vaishali Vasudeva, J.K. Garg, Ruchi Badola, Syed Ainul Hussain

## Abstract

Damming and diverting river water alters the channel characteristics and natural flow regime. The change in biotic and abiotic factors results in dissimilar habitat conditions upstream and downstream of the barrage. Given the habitat dissimilarity and therefore resource availability, we hypothesized the dissimilarity in waterbird abundance and species diversity in the river habitat upstream and downstream of the barrage. The study was conducted on a 24 km stretch of Ganga River at Narora, Uttar Pradesh, India. This stretch overlaps with a Ramsar site as well as an Important Bird and Biodiversity Area (IBA). Bird sampling was done using transect count method for summer and winter season in 2017. The abundance was studied in relation to three habitat variables *viz*. channel depth, channel width and anthropogenic disturbances using Generalized Linear Model. A total of 140 species of birds were recorded. There was statistically significant difference in the abundance of waterbirds between upstream and downstream stretch in winter season (Mann-Whitney U, *p* < 0.05). During winters, migratory waterbirds especially ducks and geese occupied the deep water upstream of barrage, while the downstream was mostly occupied by terns and cormorants. Waterbird species diversity was higher in downstream during winters (Effective Number of Species=28; Shannon’s Index= 3.35) and higher in upstream during summer (Effective Number of Species=25, Shannon’s Index=2.87). Channel width, channel depth and anthropogenic disturbance influenced waterbird abundance in both the seasons (Generalized Linear Model; *p* < 0.05). The influence of channel depth varied with season for the upstream stretch.

## Introduction

Our dependence on the surface freshwater sources is evident from the fact that “over 50% of the world’s population lives closer than 3 km to a surface freshwater body, and only 10% of the population lives further than 10 km away” [1]. Rivers around the world are being heavily modified by dams, weirs and barrages to cater for irrigation, electricity generation and drinking water requirements. These structures lead to changes in morphology, natural flow regime, flooding pattern and sedimentation in the river channel. Although, the extent to which rivers are affected is decided by factors such as dam location, size of reservoir and water residence time [2]. Nonetheless, these structures store and maintain a certain volume of water in the upstream which results in partially lentic conditions within a natural lotic system [2-6]. Alteration of environmental conditions along the regulated rivers affects the riverine biodiversity such as benthic fauna and fishes, negatively [6-10]. Besides the changes in environmental conditions, the reservoir operation and release of water in the downstream is often unpredictable being dependent upon upstream discharge. This regulation disrupts the natural flooding pattern of the river, affecting the microhabitats in the downstream and reproductive success of several ground nesting waterbirds.

Narora barrage was built on Ganga River in Northern India to supply water for irrigation through Lower Ganga Canal and Parallel Lower Ganga Canal and later to provide cooling water to the Narora Atomic Power Station. Upstream of Narora barrage, there are two water diverting structures at Haridwar and Bijnor, respectively. Due to subsequent diversion, there is a reduction of almost 85% in water flow, as Ganga river leaves Narora Barrage (4096 cusecs) compared to the water flow upstream of Haridwar barrage (30527 cusecs) [11]. This causes significant reduction in river flow downstream of Narora barrage. The barrage has a functional fish ladder that partially allows movement of fishes across the barrage but sedimentation and regulated flow have modified the natural river channel morphology. The barrage has created reservoir like conditions in the upstream with a greater water depth and partly confined channel due to hard and muddy banks. In contrast, the downstream stretch of river has highly braided and unconfined channel and more sand bars and point bars. The vegetation along the bank is sparse. Part of the river stretch being a Ramsar Site, regulates heavy fishing activities, but the unregulated stretch is disturbed with fishing nets.

Birds are conspicuous and inhabit wide variety of habitats. Changes in their population density, abundance or distribution often indicate changes in environmental conditions [12-14]. These changes can be the modification of their habitat, scarce food resources or absence of safe nesting sites. However, higher abundance or density of birds in a habitat does not indicate its higher quality [15]. Habitat selection is a behavioural response of selection of a set of specific physical environmental conditions that influence the survival of the individuals of a species [16]. Nonetheless, it has been observed that the abundance and diversity of waterbirds in a wetland, for example, is influenced by a number of drivers such as water depth, water level fluctuations vegetation type, topography, wetland size and wetland connectivity [17-21]. Water depth of a wetland and the bird body size, length of the neck and legs of bird, together with their foraging nature determines the use of specific wetland areas by birds [18, 22-23]. The variability in habitat preferences, diet and behaviour has been observed across feeding guilds and across species within a guild [24]. Birds also utilise different vertical sections of the wetland like wetland soils, or submersed and emergent plants in water column depending upon their food preferences [25].

Similarly, on a landscape scale in a riverine system, bird abundance is impacted by presence of seasonal water pools, water level, vegetation type and surface area of water [26-27]. Season is an important secondary driver that brings about change in food resources, water level and flow, water availability and affects both the species abundance and distribution of waterbirds [19-20, 28-30]. Besides natural environmental factors, birds respond to human presence and activities in positive or negative manner. It may create more foraging ground (e.g. reservoir, crop fields) or may disturb the nesting sites (e.g. fishing, construction) [28]. But largely, disturbance has been known to affect the level of habitat use and their abundance [31, 32].

Given the close association between bird morphology and habitat selection and observed dissimilarity in habitat in the river stretch upstream of barrage (henceforth USB) and downstream of barrage (henceforth DSB), we hypothesized that in the available habitat, presence of birds (habitat use) in either stretches is governed by their food habits and body size and therefore the species diversity and abundance of waterbirds is different in the two stretches. We also investigated the significance of habitat characteristics which have been modified due to construction of barrage *viz*., channel depth and channel width and the effect of anthropogenic disturbance in influencing waterbird abundance.

## Methods

### Study area

River Ganga originates from Gangotri Glacier in western Himalayas and falls into Bay of Bengal. During its course in India, it flows through five Indian states. This study was conducted in the Upper Ganga stretch at Narora, Uttar Pradesh. The study stretch lies between Karnwas (28°16’ N, 78°19’ E) and Seensai (28°06’ N, 78°28’ E) (Fig 1). The study stretch overlaps with a Ramsar site (Brijghat to Narora; 265.9 km^2^) which was declared in 2005 and also an IBA (Karnwas to Ramghat; 12,700 ha). Narora has a warm and sub-tropical climate and experiences three seasons: summer (April-June; average temp= 33.7° C), monsoon (July-September) and winter (November-March; average temperature= 14.5° C). The average annual rainfall is 902 mm. Water quality of river Ganga at Narora is good for aqua fauna, with an average dissolved oxygen (DO) of 8.2 mg/L (2006-2012), while Biological Oxygen Demand (BOD) levels show a decreasing trend from 4.0 mg/L (1999) to 2.8 mg/L (2006) [11]. The water level in the upstream of barrage is maintained at 179.07 m. The mean annual downstream gauge is 175.75 ± 1.01 m. Parallel Lower Ganga Canal receives water throughout the year with mean monthly discharge (± standard deviation *SD*) of 1704.82 ± 51.80 m^3^/s, while Lower Ganga Canal receives water only in four months (July to October) with mean monthly discharge of 110.00 ± 62.37 m^3^/s (Uttar Pradesh Irrigation Department). Throughout the study stretch, the banks are sandy with agriculture as major activity. Thousands of pilgrims visit the river banks during various Hindu festivals to perform religious and spiritual ceremonies.

**Figure 1:**
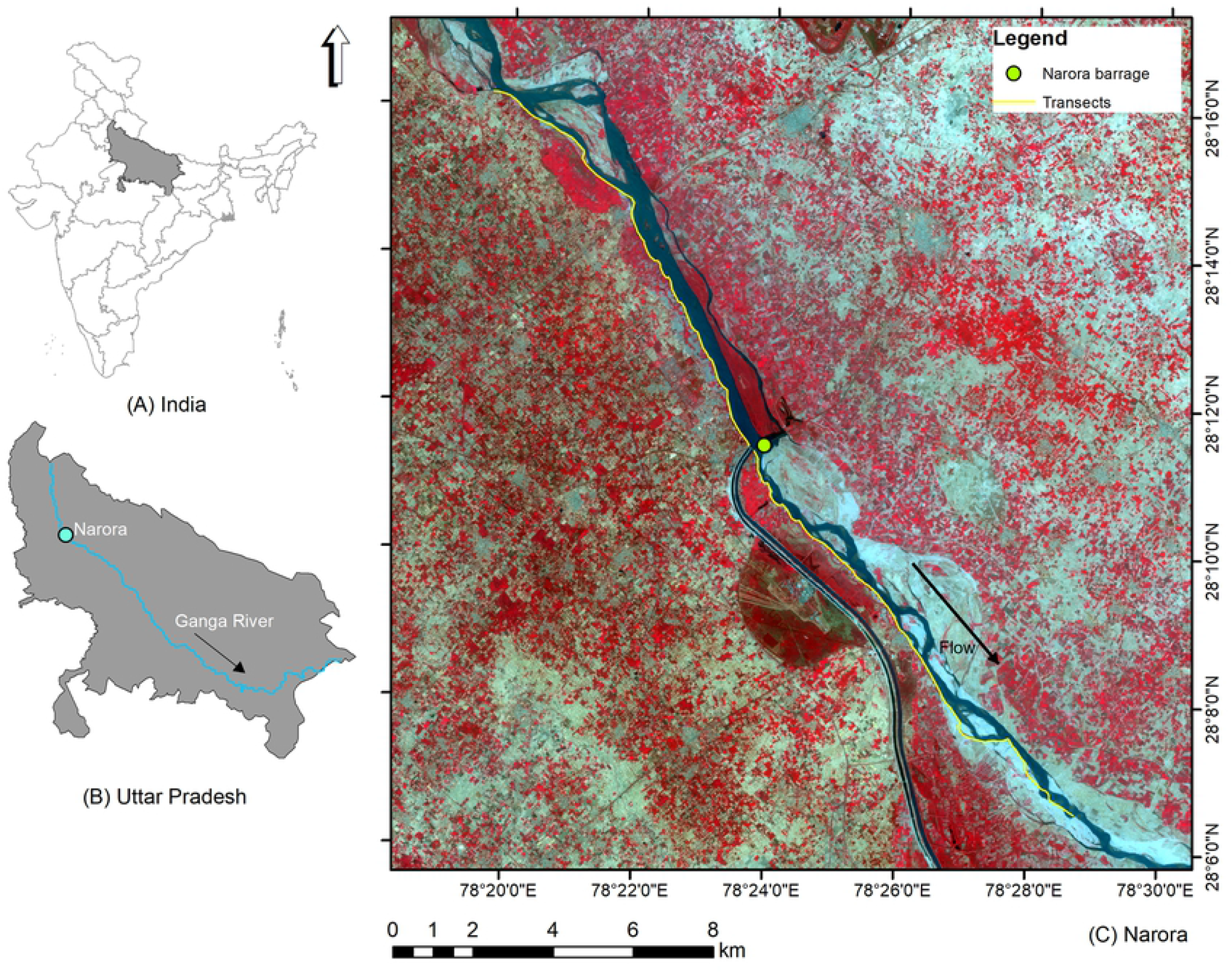
Location of study site. (A) shows location of Uttar Pradesh in India, (B) shows location of study site in Uttar Pradesh and (C) shows transects walked along the river and the location of barrage. Basemap used in (C) is the Standard False Color Composite of Sentinel-2 MSI bands (RGB: B8, B4, B3) for 27 May 2017.

Upper Ganga Ramsar site (a stretch of about 60 km from Brijghat to Narora) which also includes Narora reservoir, is an important habitat for the endangered Gangetic Dolphin *Platanista gangetica*, otters, 12 species of freshwater turtles and Mugger *Crocodylus palustris* and Gharial *Gavialis gangeticus* [33-34]. Apart from attracting thousands of migratory waterbirds every year, River Tern *Sterna aurentia*, Black-bellied Tern *Sterna acuticauda*, River Lapwing *Vanellus duvaucelli*, Sarus Crane *Grus antigone* and Indian Skimmer *Rynchops albicollis* are the important resident breeding birds in this region [35].

### Bird sampling

Ground sampling of birds was done using transect count method [36] covering a stretch of 24 km (12 km upstream and 12 km downstream of barrage). A preliminary survey was done in January 2017 and transect length and walking speed were recorded through the entire stretch (24 km). Based on the observations, the entire stretch was divided into four transects (6 km each), parallel to the river and without overlap. The starting point was Karnwas (upstream of barrage) and end point was Seensai (downstream of barrage) (Fig 1).

The bird sampling was done in the month of February 2017 for winter season and in the month of May 2017 for summer season. Five replications of each transect were made in both the seasons, making total field effort equal to 242 km for the study. Birds were continually recorded from the river bank while walking the transect on foot at a speed of 1 to 2 km/h on the right bank of the river. Birds sighted in the main channel, side channels, water pools, islands, sand bars and a strip of 10 m on the bank were recorded. The channels bifurcated only on the right bank of the river were surveyed. The birds were recorded during early morning (30 minutes after sunrise to 11 am) which corresponds to high bird activity. Birds were identified based on sightings only. For each sighting, GPS location (using hand-held GPS-Garmin 12XL), time and habitat was recorded. Only birds flying from the opposite direction parallel to river were counted and those flying across the river were excluded to avoid double counting. Observations were made for all the birds but only the count of waterbirds and water-associated birds were used in analysis. Opportunistic sightings were included in the checklist but not used in the analysis. Field Guide by [37] was used for identification. Based on food habits, food acquiring strategies and body size, we grouped the waterbirds and water-associated bird species into seven guilds (S2 Table 1).

### Habitat variables

Channel depth, channel width and anthropogenic disturbance were chosen as the appropriate primary habitat variables that signify the contrasting habitat conditions in the two stretches. Flow variations were not included as they are minimum during the period January to May, and is considered a period of lean flow for this stretch. Channel depth was measured with the help of Garmin Fishfinder 160C. Mid channel depth measurements were taken at a distance of every 100 m in the middle of river channel. Channel width was measured with the help of rangefinder (Bushnell Scout 1000 ARC Laser 5x; inclinometer accuracy= ± 0.1 degree) at a distance of every 100 m for the entire stretch. For analysis, mean depth and mean width were computed for each 1-km stretch of the river (Fig 2). Fourteen major categories of anthropogenic disturbances were identified in the study area based on observations from all ten complete surveys (S1 Table). Each category was scored on a scale of 0 to 3 depending upon intensity of disturbance for waterbirds. The scores 0 to 3 signify absence, low disturbance, moderate disturbance and high disturbance in the order. Low and moderate categories were assigned relative to the highest intensity of disturbance observed in the entire stretch and based on the observations made during all the ten visits. For each one kilometer section of the river, the scores corresponding to each category were added. Therefore, minimum possible score is zero and maximum possible score is 42.

**Figure 2:**
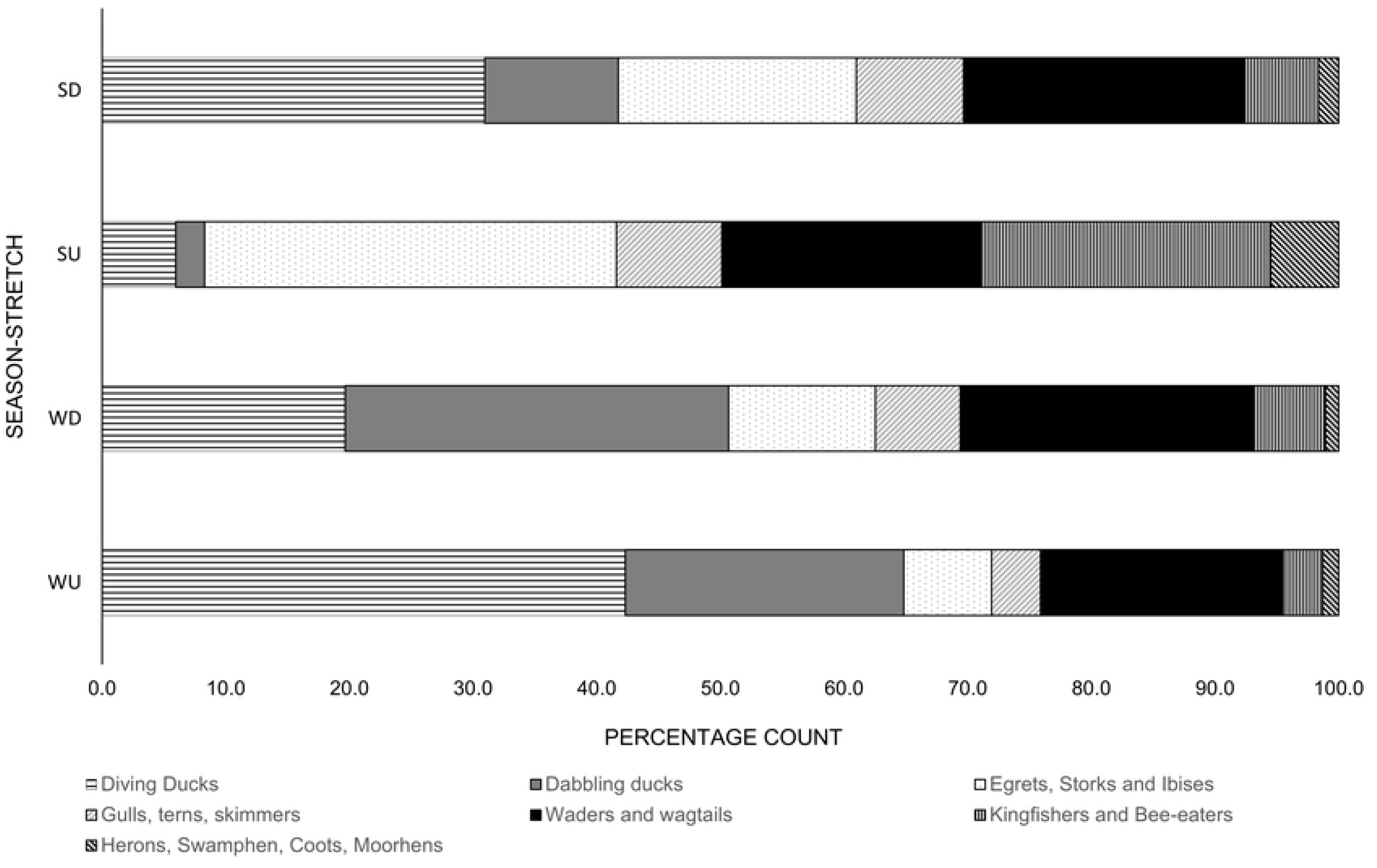
Maximum bird count for winter and summer season and habitat variables for each one km section.

### Statistical analysis

The bird count for each species from five replicates was used to compute maximum count (mode) for that species for a particular season. Maximum count was also computed for each guild considered in this study. Simpson’s Index (SI) [38], Shannon’s Diversity Index (SDI) [39], Effective Number of Species (ENS) [40] were computed to compare the species diversity upstream and downstream of the barrage. Data was checked for normality using Shapiro Wilk’s test at 95% confidence interval. Using R package ‘Flexplot’ [41] data distribution was visualized. We used Mann-Whitney U test to compare the mean between the abundance of birds in the upstream stretch and the downstream stretch. Abundance was taken as dependent variable and stretch type was taken as independent variable. Generalized Linear Model using R function ‘glm’ was used to study the influence of the three habitat variables, i.e. channel depth, channel width and anthropogenic disturbances on waterbird abundance. All analyses were carried out in R environment (R version 4.0.2, RStudio version 1.3.1073) [42].

## Results

### Overall diversity

A total of 140 species of birds were recorded within a 24 km stretch (S2 Table). These species belong to 15 orders and 42 families. Out of the 140 species, 70 species (50 %) were of waterbirds, 13 species (9%) were of water-associated birds and 57 species (40 %) of terrestrial birds. Of the total bird species observed, 39 species were resident (R), 19 were migrants (M), 21 were resident migrants (RM), 59 were resident but are subject to local movements depending upon season, availability of food and water conditions (RLM) [43]. Anatidae was the richest family of waterbirds inhabiting the river. Majority of the birds were carnivore (62 species) or omnivore (61 species) (S2 Table). During the study two endangered (EN) species: Black-bellied Tern *Sterna acuticauda* and Egyptian Vulture *Neophron percnopterus*; eleven near-threatened (NT): Ferruginous Duck *Aythya nyroca*, Painted Stork *Mycteria leucocephala*, Black-necked Stork *Ephippiorhynchus asiaticus*, Darter *Anhinga melanogaster*, Black-headed Ibis *Threskiornis melancocephalus*, Red-necked Falcon *Falco chicquera*, Great Thick-knee *Esacus neglectus*, River Tern *Sterna aurentia*, Black-tailed Godwit *Limosa limosa*, Eurasian Curlew *Numenius arquata* and River Lapwing *Vanellus duvaucelli*; and four vulnerable (VU) species: Indian Skimmer *Rynchops albicollis*, Sarus Crane *Grus antigone*, Common Pochard *Aythya ferina* and Woolly-necked Stork *Ciconia episcopus*, were observed (S2 Table). Among the breeding birds of Ganga, presence of Sarus Crane *Grus antigone*, Indian Skimmer *Rynchops albicollis*, River Lapwing *Vanellus duvaucelli*, Small Pratincole *Glareola lactea* and River Tern *Sterna aurentia* were recorded but no nesting sites were observed.

### Species richness, abundance and diversity of waterbirds and water-associated birds

In winter season, 63 species of waterbirds and water-associated birds were observed in USB and 58 species in the DSB. In summer season, 45 species were observed in USB and 44 species in DSB. Eleven species including Purple Swamphen *Porphyrio porphyrio*, Common Pochard *Aythya ferina* were exclusively observed in the USB and 13 species including Little-ringed Plover *Charadrius dubrius*, Small Pratincole *Glareola lactea* were exclusively observed in the DSB, while 60 species were common to both stretches in both the seasons (S2 Table).

A higher waterbird abundance was observed in winter season as migratory waterfowl use this site for resting and foraging from late November to early March. During winter season, 2328 birds (maximum count) were recorded in the USB and 1328 in DSB. During summer, 558 birds were recorded in the USB and 1003 birds were recorded in the DSB (Table 1). Overall abundance declined as winter migrants left the site. It was observed that number of ducks and geese declined when season changed from winter to summer but abundance of egrets, storks and ibises increased (Fig 3). Additionally, in winter, the percentage count of diving ducks was higher in the USB and in summer, it was higher in the DSB. Waders and dabbling ducks were more abundant in the DSB during summers as compared to upstream (Fig 3).

**Table 1:** Species richness, diversity indices and abundance of waterbirds and water-associated birds.

| Stretch | Season | Species<br>Richness | Simpson<br>index | Shannon's<br>Diversity Index | Effective<br>number<br>of species | Abundance<br>(Maximum<br>count) |
| --- | --- | --- | --- | --- | --- | --- |
| Upstream | Winter | 63 | 0.87 | 2.82 | 17 | 2328 |
|  | Summer | 45 | 0.95 | 3.35 | 28 | 558 |
| Downstream | Winter | 58 | 0.94 | 3.22 | 25 | 1328 |
|  | Summer | 44 | 0.91 | 2.87 | 18 | 1003 |

**Figure 3:**
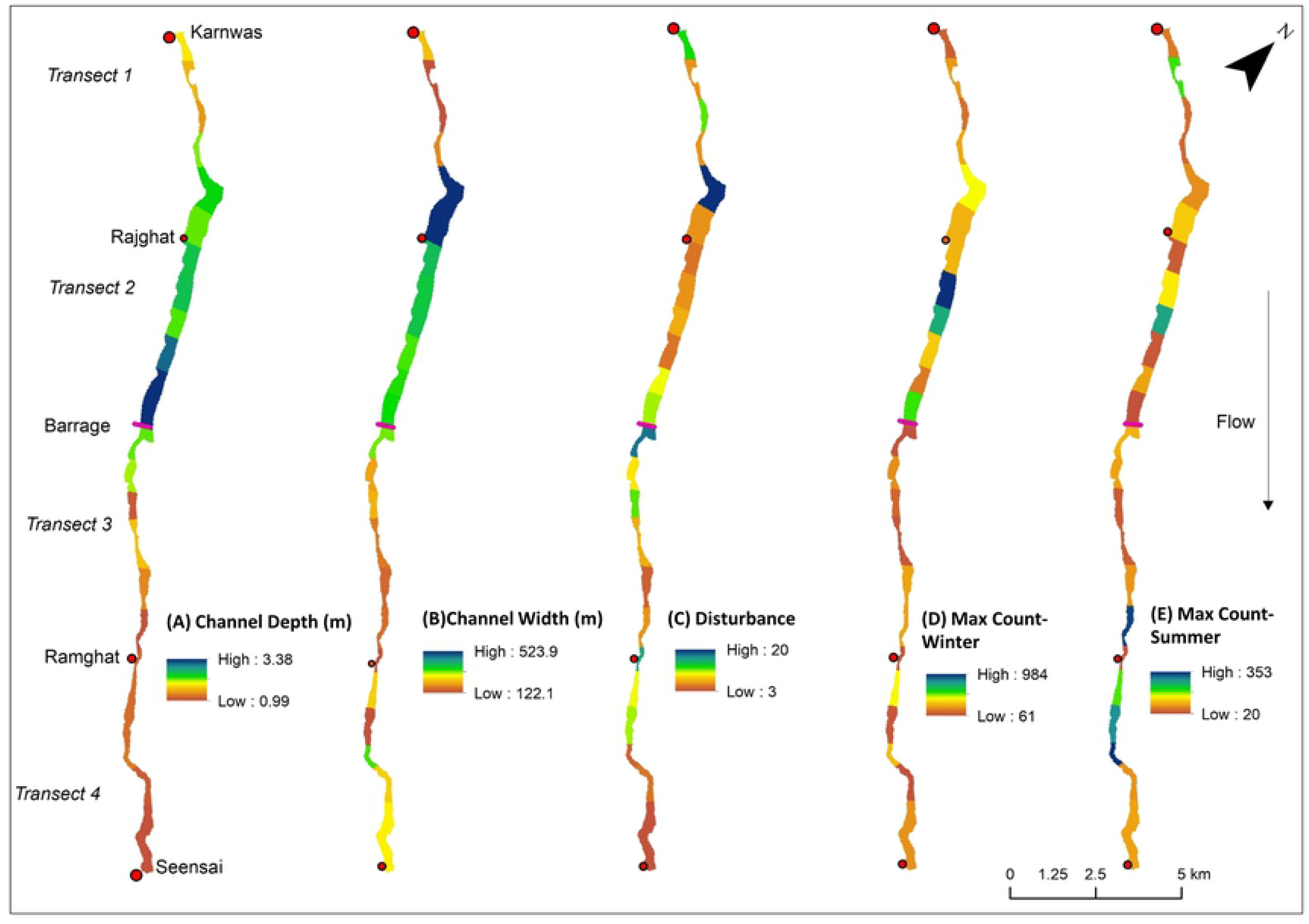
Percentage count of different waterbird groups in each season and stretch of river. WU=Winter-Upstream, WD=Winter-Downstream, SU=Summer-Upstream, SD=Summer-Downstream.

Species diversity as indicated by diversity indices, was greater in DSB in winters and in USB during summer season. In winter season, SI was higher for DSB stretch (0.94 for DSB and 0.87 for USB). Similarly, SDI and ENS were higher for DSB (SDI=3.22, ENS=25) than the USB (SDI= 2.82, ENS= 17) in winters. During summer, the USB stretch was observed to have greater diversity (SI=0.95, SDI=3.35, ENS=28) than DSB stretch (SI=0.91, SDI=2.87, ENS=18) (Table 1).

### Waterbird abundance and habitat variables

We found statistically significant difference in waterbird abundance in USB and DSB of the barrage during winter season, as indicated by Mann Whitney-U test (*p* < 0.05) where USB stretch had higher abundance of waterbirds (Table 2). The abundance was not statistically significant in summer season (*p* > 0.05) (Table 2).

**Table 2:** Test parameters and output for Mann-Whitney U test for difference in mean bird abundance in the USB and DSB.

| Season | Stretch | Mean Rank | Sum of Ranks | Mann-Whitney U | z | Significance |
| --- | --- | --- | --- | --- | --- | --- |
| Winter | Upstream | 8 | 40 | 0.0 | -2.61 | 0.009 |
|  | Downstream | 3 | 15 |  |  |  |
| Summer | Upstream | 5.6 | 28 | 12.0 | -0.104 | 0.917 |
|  | Downstream | 5.4 | 27 |  |  |  |

The influence of channel width, channel depth and disturbance on waterbird abundance was statistically significant in both the seasons (*p* < 0.05) (Table 3). Disturbance had negative influence and channel width had a positive influence on waterbird abundance in both the seasons. Channel depth had a positive influence on abundance in winter season (only in the upstream) and negative in summer season (Table 3).

**Table 3:**
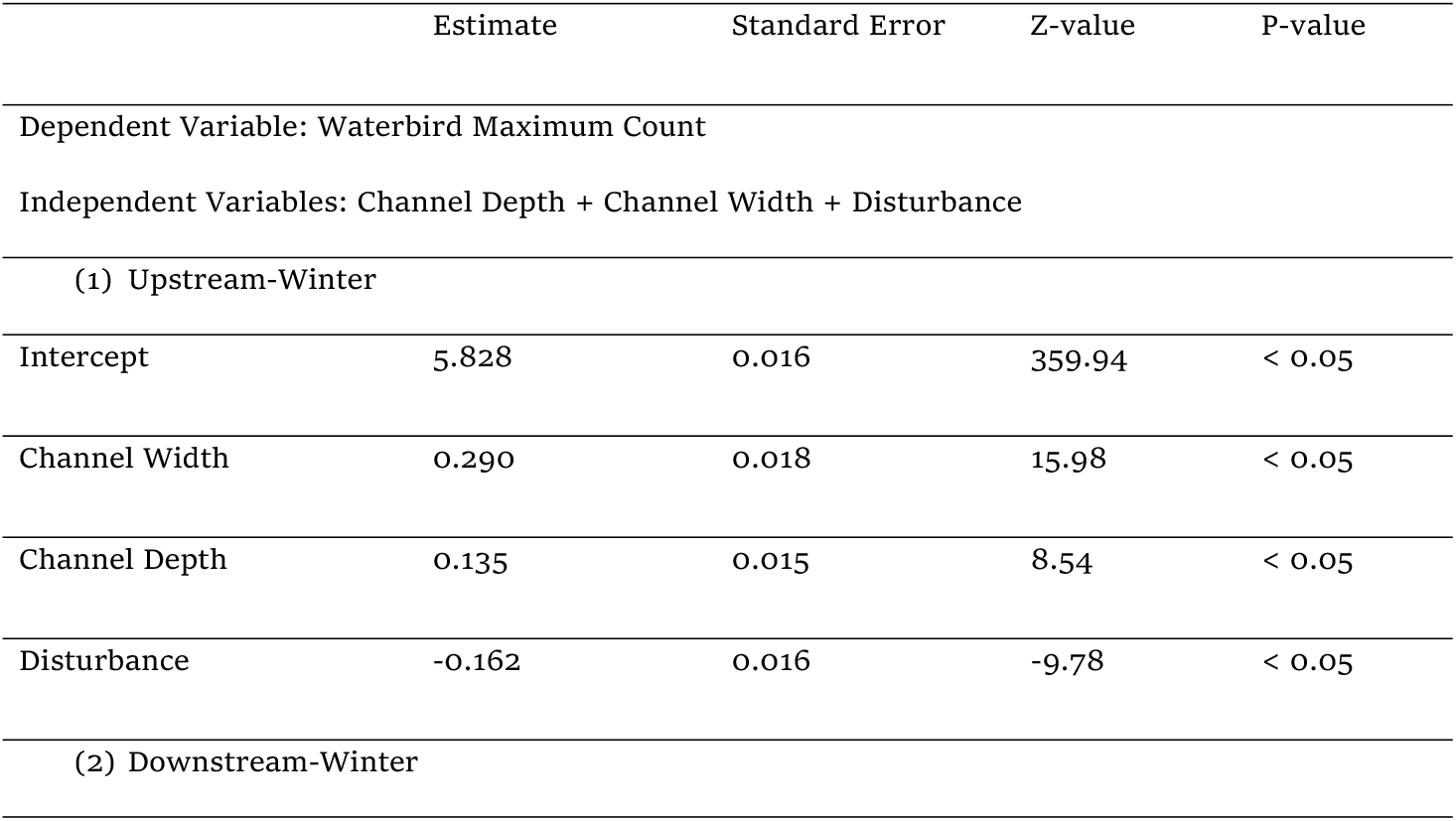
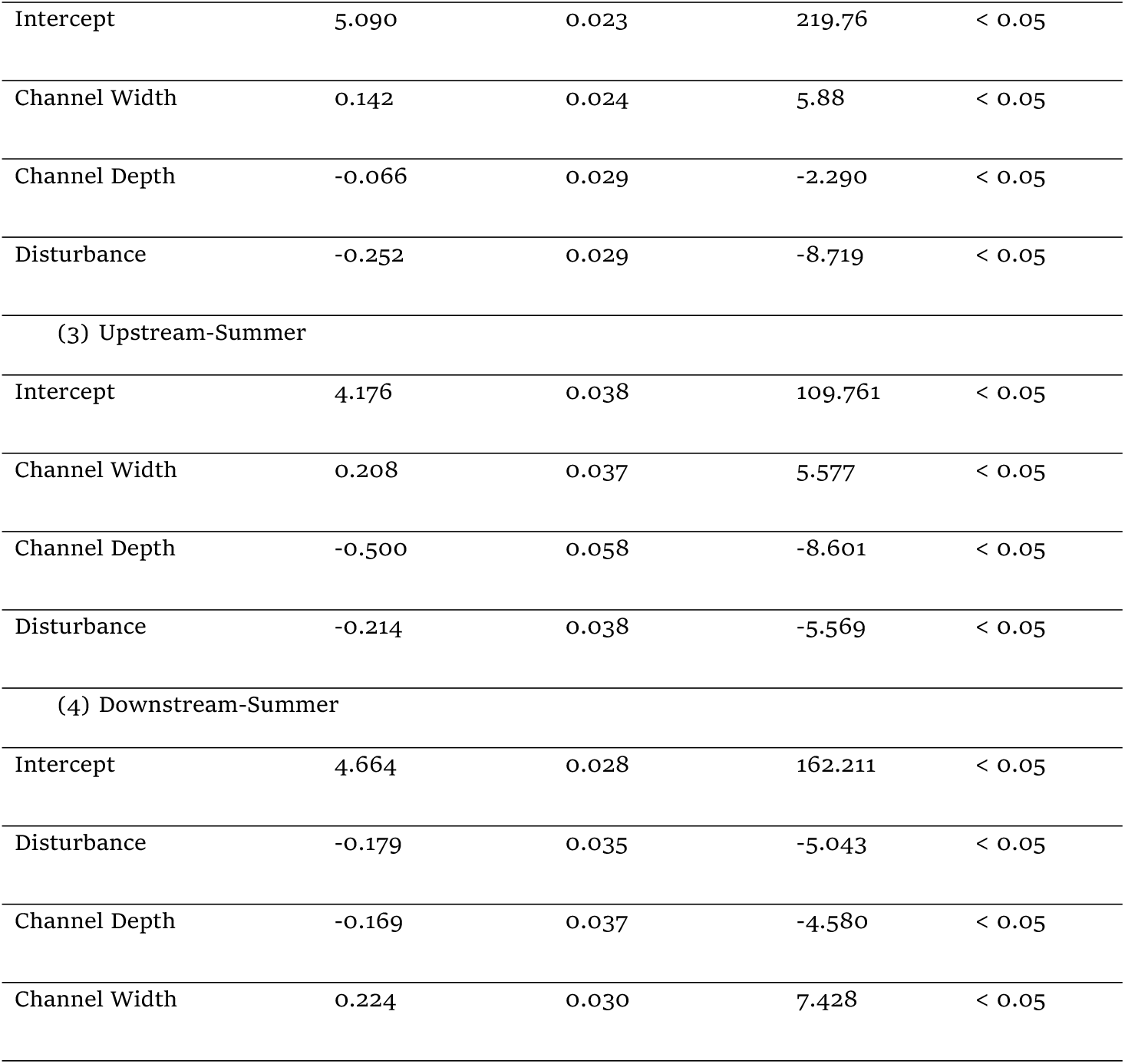
Output of different generalized linear models for influence of habitat variables on waterbird abundance in USB and DSB.

|  | Estimate | Standard Error | Z-value | P-value |
| --- | --- | --- | --- | --- |
| Dependent Variable: Waterbird Maximum Count |  |  |  |  |
| Independent Variables: Channel Depth + Channel Width + Disturbance |  |  |  |  |
| (1) Upstream-Winter |  |  |  |  |
| Intercept | 5.828 | 0.016 | 359.94 | < 0.05 |
| Channel Width | 0.290 | 0.018 | 15.98 | < 0.05 |
| Channel Depth | 0.135 | 0.015 | 8.54 | < 0.05 |
| Disturbance | -0.162 | 0.016 | -9.78 | < 0.05 |
| (2) Downstream-Winter |  |  |  |  |
| Intercept | 5.090 | 0.023 | 219.76 | < 0.05 |
| Channel Width | 0.142 | 0.024 | 5.88 | < 0.05 |
| Channel Depth | -0.066 | 0.029 | -2.290 | < 0.05 |
| Disturbance | -0.252 | 0.029 | -8.719 | < 0.05 |
| (3) Upstream-Summer |  |  |  |  |
| Intercept | 4.176 | 0.038 | 109.761 | < 0.05 |
| Channel Width | 0.208 | 0.037 | 5.577 | < 0.05 |
| Channel Depth | -0.500 | 0.058 | -8.601 | < 0.05 |
| Disturbance | -0.214 | 0.038 | -5.569 | < 0.05 |
| (4) Downstream-Summer |  |  |  |  |
| Intercept | 4.664 | 0.028 | 162.211 | < 0.05 |
| Disturbance | -0.179 | 0.035 | -5.043 | < 0.05 |
| Channel Depth | -0.169 | 0.037 | -4.580 | < 0.05 |
| Channel Width | 0.224 | 0.030 | 7.428 | < 0.05 |

Analysis of the influence of habitat variables for different guilds revealed that channel depth and width are significant predictors of abundance for all the guilds (Table 4). Disturbance had statistically significant influence on abundance of all the guilds except for G5 (Table 4). Stretch (USB or DSB) was found to be significant for all guilds except for G7 indicating that the abundance of water-associated birds is not influenced by the stretch type. Season did not influence the abundance of guilds G3, G6 and G7 (Table 4).

**Table 4:** Output of different generalized linear models for influence of habitat variables in the on waterbird abundance of each guild. (W) and (U) indicate winter and upstream respectively.

|  | Variable | Estimate | Standard Error | Z-Value | P value |
| --- | --- | --- | --- | --- | --- |
| All waterbirds and water-associated birds | Intercept | 4.132 | 0.046 | 88.888 | <0.05 |
|  | Depth | 0.053 | 0.026 | 2.007 | 0.04 |
|  | Width | 0.002 | 0.000 | 18.908 | <0.05 |
|  | Disturbance | -0.045 | 0.002 | -17.437 | <0.05 |
|  | Season (W) | 1.061 | 0.024 | 42.993 | <0.05 |
|  | Stretch (U) | 0.158 | 0.031 | 5.087 | <0.05 |
| G1: Diving ducks, Cormorants and Grebes | Intercept | 0.468 | 0.099 | 4.704 | <0.05 |
|  | Depth | 1.023 | 0.037 | 27.102 | <0.05 |
|  | Width | 0.005 | 0.000 | 22.489 | <0.05 |
|  | Disturbance | -0.070 | 0.005 | -14.193 | <0.05 |
|  | Season (W) | 1.562 | 0.050 | 30.762 | <0.05 |
|  | Stretch (U) | -0.773 | 0.070 | -10.918 | <0.05 |
| G2: Dabbling Ducks | Intercept | 4.118 | 0.139 | 29.488 | <0.05 |
|  | Depth | -1.314 | 0.099 | -13.183 | <0.05 |
|  | Width | 0.001 | 0.000 | 4.164 | <0.05 |
|  | Disturbance | -0.135 | 0.007 | -17.621 | <0.05 |
|  | Season (W) | 2.272 | 0.081 | 27.871 | <0.05 |
|  | Stretch (U) | 1.093 | 0.073 | 14.905 | <0.05 |
| G3: Egrets, Storks and Ibises | Intercept | 3.822 | 0.136 | 28.098 | <0.05 |
|  | Depth | -0.783 | 0.102 | -7.621 | <0.05 |
|  | Width | 0.002 | 0.000 | 7.580 | <0.05 |
|  | Disturbance | 0.042 | 0.007 | -5.883 | <0.05 |
|  | Season (W) | 0.010 | 0.060 | 0.181 | 0.856 |
|  | Stretch (U) | 0.467 | 0.087 | 5.320 | <0.05 |
| G4: Gulls, Terns and Skimmers | Intercept | 1.297 | 0.207 | 6.260 | <0.05 |
|  | Depth | -0.503 | 0.139 | -3.600 | <0.05 |
|  | Width | 0.005 | 0.000 | 10.845 | <0.05 |
|  | Disturbance | 0.027 | 0.008 | 3.231 | 0.001 |
|  | Season (W) | 0.492 | 0.092 | 5.352 | <0.05 |
|  | Stretch (U) | -0.411 | 0.141 | -2.900 | 0.003 |
| G5: Waders and Wagtails | Intercept | 2.991 | 0.101 | 29.521 | <0.05 |
|  | Depth | -0.429 | 0.067 | -6.389 | <0.05 |
|  | Width | 0.001 | 0.000 | 7.096 | <0.05 |
|  | Disturbance | 0.005 | 0.004 | 1.071 | 0.284 |
|  | Season (W) | 1.007 | 0.052 | 19.033 | <0.05 |
|  | Stretch (U) | 0.371 | 0.064 | 5.795 | <0.05 |
| G6: Herons, Swamphen, Coots and Moorhens | Intercept | 4.189 | 0.200 | 20.899 | <0.05 |
|  | Depth | -0.908 | 0.158 | -5.726 | <0.05 |
|  | Width | -0.000 | 0.000 | -1.978 | 0.047 |
|  | Disturbance | -0.098 | 0.012 | -7.675 | <0.05 |
|  | Season (W) | -0.077 | 0.086 | -0.903 | 0.366 |
|  | Stretch (U) | 1.236 | 0.120 | 10.229 | <0.05 |
| G7: Kingfishers, Bee-eaters and Martins | Intercept | -0.374 | 0.357 | -1.047 | 0.294 |
|  | Depth | 0.906 | 0.165 | 5.492 | <0.05 |
|  | Width | 0.002 | 0.000 | 2.805 | 0.005 |
|  | Disturbance | -0.104 | 0.025 | -4.159 | <0.05 |
|  | Season (W) | 0.122 | 0.165 | 0.742 | 0.458 |
|  | Stretch (U) | -0.223 | 0.271 | -0.823 | 0.410 |

## Discussion

Previously only two studies have been conducted at this site following different method and effort. [35] recorded 120 species at Narora including Greylag goose, Common Shelduck, White-eyed Pochard and Shoveller, which were not observed during this study duration. Similarly, several pairs or individuals of Black-necked Stork *Ephippiorhynchus asiaticus* were seen in the lakes and shallow areas of the reservoir by [35]. In the present study we recorded one individual of Black-necked Stork in the DSB. [44] recorded 55 species of birds in 2007 during a survey, which was carried out from Bijnor to Narora, covering a larger stretch. They recorded species such as Greater Flamingo (*Phoenicopterus ruber*), Common Teal (*Anas crecca*), Pintail (*Anas acuta*), Northern Shoveller (*Anas clypeata*) which were not recorded during our sampling. This could be attributed to difference in survey effort and extent in the two studies but certainly highlights the need for long term studies in order to systematically monitor the population of birds and their habitat.

Water depth is an important factor that can differentiate between waterbird habitats [23]. During winters, river depth is the positive influencing factor in the USB. This positive influence of depth in USB was due to presence of reservoir and increased surface area of water which is preferred by diving ducks and dabbling birds [32]. It was observed to be a preferred habitat by Common Pochard, which was only observed in USB. In summer, we observed a negative influence of water depth on waterbird abundance. This observation can be attributed to change in species composition with change in season, where diving and dabbling ducks were dominating in the winters but waders, herons and kingfishers were more abundant in the summer season and occupied shallow water habitats (Fig 2). In DSB, where water depth is lower than USB, a mosaic of habitat due to exposed point bars and islands is available which provided additional habitat for shallow water preferring birds [45]. Waders like small pratincole and little ringed plover were exclusively found in the downstream stretch, as they prefer areas with lowest vegetation cover [46].

Similarly, we found that channel width positively influences the abundance of diving ducks (G1) by increasing the area of available habitat and disturbance negatively affects the abundance of waterbirds, irrespective of site and season. Birds avoided sites which were intensively used by humans but some of the disturbances like fishing nets, agriculture and religious offerings attracted birds due to food availability, for example in case of Gulls, terns, skimmers and Egrets, storks & ibises. The consequent increase in the abundance was not reflected in the results as it could have been masked by simultaneous increase in river width. Absence of river bed farming in the downstream stretch provided sandy bars and islands to be used by large flocks of cormorants and egrets. Thus, absence of anthropogenic activities is an important factor influencing the use of habitat by birds.

Other factors that could have affected the abundance and diversity of birds are habitat diversity, food availability and daily water flow fluctuations. For example, as observed during the present study and by [17] that when the islands were submerged due to flow variations, more birds were observed on the shores and banks. The guild-wise response of waterbird abundance to habitat variables cannot be explained considering the habitat variables alone. Further studies are suggested to incorporate additional variables for a better understanding of habitat use and selection by waterbirds in different seasons. This region provides an important habitat to both migratory and resident birds. The reservoir created by barrage supports migratory waterfowl by providing suitable habitat but unfavorable impacts on other aquatic taxa and sediment retention are important issues. The water released by the barrage during breeding season of birds must be regulated, taking into account the submergence of islands and point bars. Other activities like river-bed farming and fishing with nets must also be regulated to make the site suitable for nesting of breeding birds of Ganga River.

This study was a preliminary attempt to understand the difference in waterbird abundance and diversity in the partially lacustrine and lotic habitat conditions created by a water diverting structure on Ganga River. Also this study is the first intensive study that covers part of both the Upper Ganga Ramsar Site and the Narora IBA. It can be concluded that the two stretches upstream and downstream of barrage have dissimilar abundance and species diversity of waterbirds that varies with season. The influence of channel depth, channel width, and anthropogenic disturbance on waterbird abundance was statistically significant in both seasons.

## Acknowledgements

We thank the Director and the Dean, Wildlife Institute of India for extending us this opportunity to carry out this piece of research. We thank Ms Monika Mehralu, Mr Gaura Chandra Das and Mr Aftab Usmani for assisting in field data collection and the authorities of Narora Atomic Power Station for the logistic support on the field.

## Supporting information captions

**S1 Table:** Disturbance categories identified in quantification of disturbance for the waterbirds at the study site and their description.

**S2 Table:** Checklist of birds observed at Narora.

